# Aberrant Dynamic Functional Connectivity of Default Mode Network in Schizophrenia and Links to Symptom Severity

**DOI:** 10.1101/2021.01.03.425152

**Authors:** Mohammad S.E. Sendi, Elaheh Zendehrouh, Charles A. Ellis, Zhijia Liang, Zening Fu, Daniel H. Mathalon, Judith M. Ford, Adrian Preda, Theo G. M. van Erp, Robyn. L Miller, Godfrey D. Pearlson, Jessica A. Turner, Vince D. Calhoun

## Abstract

**Background:** Schizophrenia affects around 1% of the global population. Functional connectivity extracted from resting-state functional magnetic resonance imaging (rs-fMRI) has previously been used to study schizophrenia and has great potential to provide novel insights into the disorder. Some studies have shown abnormal functional connectivity in the default mode network of individuals with schizophrenia, and more recent studies have shown abnormal dynamic functional connectivity (dFC) in individuals with schizophrenia. However, DMN dFC and the link between abnormal DMN dFC and symptom severity have not been well characterized.

**Method:** Resting-state fMRI data from subjects with schizophrenia (SZ) and healthy controls (HC) across two datasets were analyzed independently. We captured seven maximally independent subnodes in the DMN by applying group independent component analysis and estimated dFC between subnode time courses using a sliding window approach. A clustering method separated the dFCs into five reoccurring brain states. A feature selection method modeled the difference between SZs and HCs using the state-specific FC features. Finally, we used the transition probability of a hidden Markov model to characterize the link between symptom severity and dFC in SZ subjects.

**Results:** We found decreases in the connectivity of the anterior cingulate cortex (ACC) and increases in the connectivity between the precuneus (PCu) and the posterior cingulate cortex (PCC) (i.e., PCu/PCC) of SZ subjects. In SZ, the transition probability from a state with weaker PCu/PCC and stronger ACC connectivity to a state with stronger PCu/PCC and weaker ACC connectivity increased with symptom severity.

**Conclusions:** To our knowledge, this was the first study to investigate DMN dFC and its link to schizophrenia symptom severity. We identified reproducible neural states in a data-driven manner and demonstrated that the strength of connectivity within those states differed between SZs and HCs. Additionally, we identified a relationship between SZ symptom severity and the dynamics of DMN functional connectivity. We validated our results across two datasets. These results support the potential of dFC for use as a biomarker of schizophrenia and shed new light upon the relationship between schizophrenia and DMN dynamics.

## 1 Introduction

In recent years, static functional connectivity (sFC) obtained from resting-state functional magnetic resonance imaging (rs-fMRI) time series has revealed a great deal of knowledge about brain dysconnectivity in schizophrenia (Lynall et al., 2010; Skåtun et al., 2017). Among intrinsic brain networks, the default mode network (DMN) – including the anterior cingulate cortex (ACC), posterior cingulate cortex (PCC), precuneus (PCu), medial prefrontal cortex (mPFC), ventral ACC, and the lateral/inferior parietal cortices – has been widely studied due to its putative role in external monitoring, spontaneous cognition, and autobiographical thinking (Hu et al., 2017) and due to its links to mental disorders like schizophrenia (Du et al., 2016).

In the DMN, the anterior and posterior cingulate cortices (ACC and PCC) are involved in multiple complex cognitive functions, including decision-making, empathy, emotion, socially-driven interactions, and autobiographical memory (Stevens et al., 2011; Leech and Sharp, 2014). Several studies showed a functional and structural alteration within and between the cingulate cortex and other regions that emphasized the role of this region in the pathology of schizophrenia (Wood et al., 2007; Calabrese et al., 2008; Whitfield-Gabrieli et al., 2009; Woodward et al., 2011; Yan et al., 2012; Peeters et al., 2015; Wang et al., 2015, 2017; Guo et al., 2017; Li et al., 2019). In a voxel-wise comparison between SZs and HCs, SZ individuals show a reduction of ACC gray matter (Wang et al., 2007). In addition, a reduction of ACC functional connectivity within DMN has been associated with SZ (Li et al., 2019). Regarding the PCC, a reduction of PCC gray matter volume has been found in both individuals with schizophrenia and their non-psychotic siblings (Calabrese et al., 2008). One rs-fMRI study showed higher connectivity between the PCC and PCu in SZ subjects (Whitfield-Gabrieli et al., 2009). Consistent with this, an increase in connectivity between the PCu and PCC has been reported in schizophrenia subjects and their siblings (Peeters et al., 2015). In a small sample size, lower functional connectivity of the ACC in the anterior DMN and the PCu in the posterior DMN of schizophrenia subjects exhibiting poor insight is reported (Liemburg et al., 2012).

Several studies from our group and others have previously reported a link between sFC among the ACC, PCC, and PCu and symptom severity in schizophrenia (Whitfield-Gabrieli et al., 2009; Chain et al., 2019; Hare et al., 2019). One of those studies reported a positive correlation between PCu/PCC connectivity and symptom severity as measured by the scale for the assessment of positive symptoms (SAPS) in a relatively small number of subjects (Whitfield-Gabrieli et al., 2009). A separate study showed aberrant connectivity within the DMN and also that DMN connectivity correlates with symptom severity in schizophrenia subjects (Garrity et al., 2007), and another study found a link between the ACC thickness of SZ subjects and the duration of illness and severity of psychotic symptoms (Wang et al., 2007).

All the studies mentioned above either studied the DMN as a whole or emphasized the separate role of the PCC, ACC, and PCu within the DMN and their connectivity to the pathology of schizophrenia. However, inconsistent results in the functional connectivity of the regions have been previously observed. For example, previous studies showed that SZ subjects had both an increase and a decrease in ACC connectivity within the DMN compared with HC (Li et al., 2019; Shukla et al., 2019). Although this inconsistency could, to a limited extent, be attributed to differences in disease subtypes or symptoms, we theorize that some of the heterogeneity is driven by the emphasis on sFC, which represents an average across different brain states during an unconstrained resting state.

Unlike conventional sFC, which is obtained from the correlation within an entire time series, dynamic functional connectivity (dFC) or its network analog, dynamic functional network connectivity (dFNC) refers to the connectivity between pairs of brain regions (or networks) within sub-intervals of time series (Calhoun et al., 2014). In fact, dFC research suggests that cognitive deficits and clinical symptoms associated with many psychiatric disorders not only depend on the strength of the connectivity between any brain regions but also on the variation of connectivity strength between those regions over time (Calhoun et al., 2014; Damaraju et al., 2014; Du et al., 2015; Engels et al., 2018; Vergara et al., 2018; Bhinge et al., 2019; Sanfratello et al., 2019; Schumacher et al., 2019).

The temporal feature of dFC has been reported as a plausible biomarker for identifying the fundamental mechanisms differentiating healthy individuals and schizophrenia subjects (Damaraju et al., 2014; Du et al., 2015; Rashid et al., 2016; Sanfratello et al., 2019). A previous whole-brain dynamic connectivity analysis showed that schizophrenia subjects spend less time in a highly-connected state (Damaraju et al., 2014; Sendi et al., 2020). A study from our group showed an abnormal pattern in the dFNC of the DMN by comparing state-based connectivity strength, dwell time, and the number of between-state transitions of HC and SZ subjects (Du et al., 2015). The study identified a SZ-associated pattern in the temporal dynamics of the DMN in SZ subjects by showing that they spend more time in a state with sparsely connected nodes. The study also demonstrated a state-specific spatial disruption within the DMN by showing that the central hubs of the PCC and the anterior medial prefrontal cortex are significantly impaired in SZ subjects. However, the study did not show how symptom severity is associated with the identified abnormal pattern and how dFC patterns differ between subjects with varying symptom severity. Also, in contrast with the previously mentioned study that used a seed-based approach to extract the brain network components (regions), in the current study, we used a framework called NeuroMark (Du et al., 2019). NeuroMark is a fully automated independent component analysis (ICA) framework that uses spatially constrained ICA to estimate comparable features across subjects by taking advantage of the replicated brain network templates extracted from two N∼900 normative resting fMRI data sets. We analyzed the dFC of data-driven DMN subnodes based on the NeuroMark template and showed an aberrant temporal pattern and a link between this connectivity pattern and symptom severity in schizophrenia.

To investigate the temporal dynamics of FNC within DMN subnode connectivity, we used two different datasets. A sliding window approach was used to generate dFC samples, and k-means clustering was applied to identify a set of data-driven dFC states (Calhoun et al., 2014). Further, to investigate and model the temporal changes in the dFC, we estimated the transition probability via a hidden Markov model (HMM) applied to the dFC data. In the next step, via statistical analysis on the estimated HMM features, we tested for links between schizophrenia symptom severity and abnormal DMN dFC. Finally, to investigate within-state variability across all subjects, we utilized an interpretable machine learning approach, called logistic regression with elastic net regularization (ENR), to identify the features that were most important to differentiating between SZ and HC subjects (Tibshirani, 2011). This approach can model the differences between SZ and HC individuals in the connectivity of DMN subnodes within each state. We hypothesized that the disruption of state-dependent connectivity within a shorter timescale would reveal more information about the dynamics among DMN subnodes in schizophrenia and potentially explain previous heterogeneous findings regarding these subnodes. Also, the investigation of the link between symptom severity and dFC within the three network subnodes provides additional insight into the link between functional connectivity dynamics and clinical phenomenology. The application of these methods to two distinct rs-fMRI datasets enabled the validation our findings and increased the likelihood of our results being generalizable across the broader population of individuals with schizophrenia.

## 2 Materials and Methods

### 2.1 Participants and Dataset

Data were obtained from the Mind Research Network Center of Biomedical Research Excellence (COBRE) (Aine et al., 2017) and the Functional Imaging Biomedical Informatics Research Network (FBIRN) (van Erp et al., 2015) projects. The COBRE dataset includes 89 HCs and 68 SZ subjects. The FBIRN dataset contains 151 SZ subjects and 160 HCs. The raw imaging data were collected from seven sites including the University of California, Irvine; the University of California, Los Angeles; the University of California, San Francisco; Duke University/the University of North Carolina at Chapel Hill; the University of New Mexico; the University of Iowa; and the University of Minnesota. In this study, written informed consent was obtained from all participants. Institutional review boards approved the consent process of each study site. It is worth mentioning that the COBRE subjects’ eyes were open during scanning while the FBIRN subjects’ eyes were closed. SZ patients were on a stable dose of typical, atypical, or combination antipsychotic medication for at least two months prior to data recording and had an illness duration of at least 1 year. HC and SZ individuals with a history of significant medical illness and an IQ of less than 75 were excluded from the study. In addition, those HC subjects with a current or past history of major neurological and psychiatric disorders in either themselves or a first-degree relative were excluded from this study. The demographic information for these subjects is shown in Table 1 and Supplementary Table S.1. Using a two-sample t-test, we did not observe a significant difference between the ages of the HC and SZ groups in either dataset. A diagnosis of schizophrenia was confirmed with the SCID-IV interview (First et al., 2002b), and an absence of schizophrenia diagnosis in HC was confirmed with the SCID-I/NP interview (First et al., 2002a). In addition, HCs with a first-degree relative with an Axis-I psychotic disorder diagnosis were also excluded. Symptom scores were determined based on the positive and negative syndrome scale (PANSS) (Hare et al., 2017).

**Table 1.** Demographic and clinical information of subjects

|  |  | *SZ* | *HC* | *P-value* |
| --- | --- | --- | --- | --- |
| *C* | Number | 68 | 89 | NA |
| *O* | Age | 37.79±14.44 | 38.09±11.66 | 0.52 |
| *B* | Gender(M/F) | 57/11 | 64/25 | 0.61 |
| *R* | PANSS (positive) | 15.29±5.05 | NA | NA |
| *E* | PANSS (negative) | 14.72±5.45 | NA | NA |
| *F* | Number | 151 | 160 | NA |
| *B* | Age | 38.06±11.30 | 37.04±10.68 | 0.41 |
| *I* | Gender(M/F) | 115/36 | 115/45 | 0.99 |
| *R* | PANSS (positive) | 15.32±4.92 | NA | NA |
| *N* | PANSS (negative) | 14.32±5.42 | NA | NA |

**Note:** SZ: Schizophrenia; HC: healthy control; PANSS: Positive and Negative Syndrome Scale, M: Male, F: Female, NA: not applicable; all p values have been calculated using two-sample t-test.

### 2.2 MRI Data Acquisition System

For the COBRE dataset, the MRI Images were collected on a single 3-Tesla Siemens Trio scanner with a 12-channel radiofrequency coil. High resolution T2*-weighted functional images were acquired using a gradient-echo echo-planar imaging (EPI) sequence with TE = 29 ms, TR = 2 s, flip angle = 75°, slice thickness = 3.5 mm, slice gap = 1.05 mm, field of view = 240 mm, matrix size = 64, voxel size = 3.75 × 3.75 × 4.55 mm^3^. Resting-state scans consisted of 149 volumes. Subjects were instructed to keep their eyes open during the resting-state scan and stare passively at a central cross (Aine et al., 2017). For the FBIRN dataset, six sites used 3T Siemens TIM Trio scanners, and one site used a 3T GE MR750 scanner for collecting the imaging data. All sites used the following T2*-weighted AC-PC aligned EPI sequence: TR = 2 s, TE = 30 ms, flip angle = 77°, slice gap = 1 mm, voxel size = 3.4 × 3.4 × 4 mm^3^, number of frames = 162, acquisition time = 5 min and 38 s (van Erp et al., 2015).

### 2.3 Data Processing

Statistical parametric mapping (SPM12, https://www.fil.ion.ucl.ac.uk/spm/) in the MATLAB2019 environment was used to preprocess fMRI data. The first five dummy scans were discarded before preprocessing. Slice-timing correction was performed on the fMRI data. Rigid body motion correction was then applied to account for subject head motion in SPM. Next, the imaging data underwent spatial normalization to an EPI template in the standard Montreal Neurological Institute (MNI) space and was resampled to 3×3×3 mm^3^. Finally, a Gaussian kernel was used to smooth the fMRI images using a full width at half maximum (FWHM) of 6 mm.

In each dataset, to extract reliable DMN independent components (ICs), we used the Neuromark automatic ICA pipeline within the group ICA of fMRI toolbox (GIFT, http://trendscenter.org/software/gift), which uses previously derived component maps as priors for spatially constrained ICA (Du et al., 2019). The Neuromark automatic ICA pipeline was used to extract ICs by employing previously-derived component maps as priors for spatially constrained ICA. In Neuromark, replicable components were identified by matching group-level spatial maps from two large-sample HC datasets. Components were identified as meaningful regions if they exhibited peak activations in the gray matter within the DMN. Seven DMN subnodes were identified based on an anatomical template (Tzourio-Mazoyer et al., 2002). This set of subnodes included three subnodes in the PCu, two subnodes in the ACC, and two subnodes in the PCC. These subnodes are shown in Table 2 and Figure 1 (Step 1). With seven DMN subnodes, we had twenty-one connectivity features, where each feature represented the strength of the connection between a pair of DMN subnodes.

**Table 2.** Component labels extracted using Neuromark

| Component name | Peak coordinate (mm) |  |  |
| --- | --- | --- | --- |
| (IC 32), Precuneus [PCu1] | -8.5 | -66.5 | 35.5 |
| (IC,40), Precuneus [PCu2] | -12.5 | -54.5 | 14.5 |
| (IC 23), Anterior cingulate cortex [ACC1] | -2.5 | 35.5 | 2.5 |
| (IC 71), Posterior cingulate cortex [PCC1] | -5.5 | -28.5 | 26.5 |
| (IC 17), Anterior cingulate cortex [ACC2] | -9.5 | 46.5 | -10.5 |
| (IC 51), Precuneus [PCu3] | -0.5 | -48.5 | 49.5 |
| (IC 94), Posterior cingulate cortex ([PCC2] | -2.5 | 54.5 | 31.5 |

**Figure 1.**
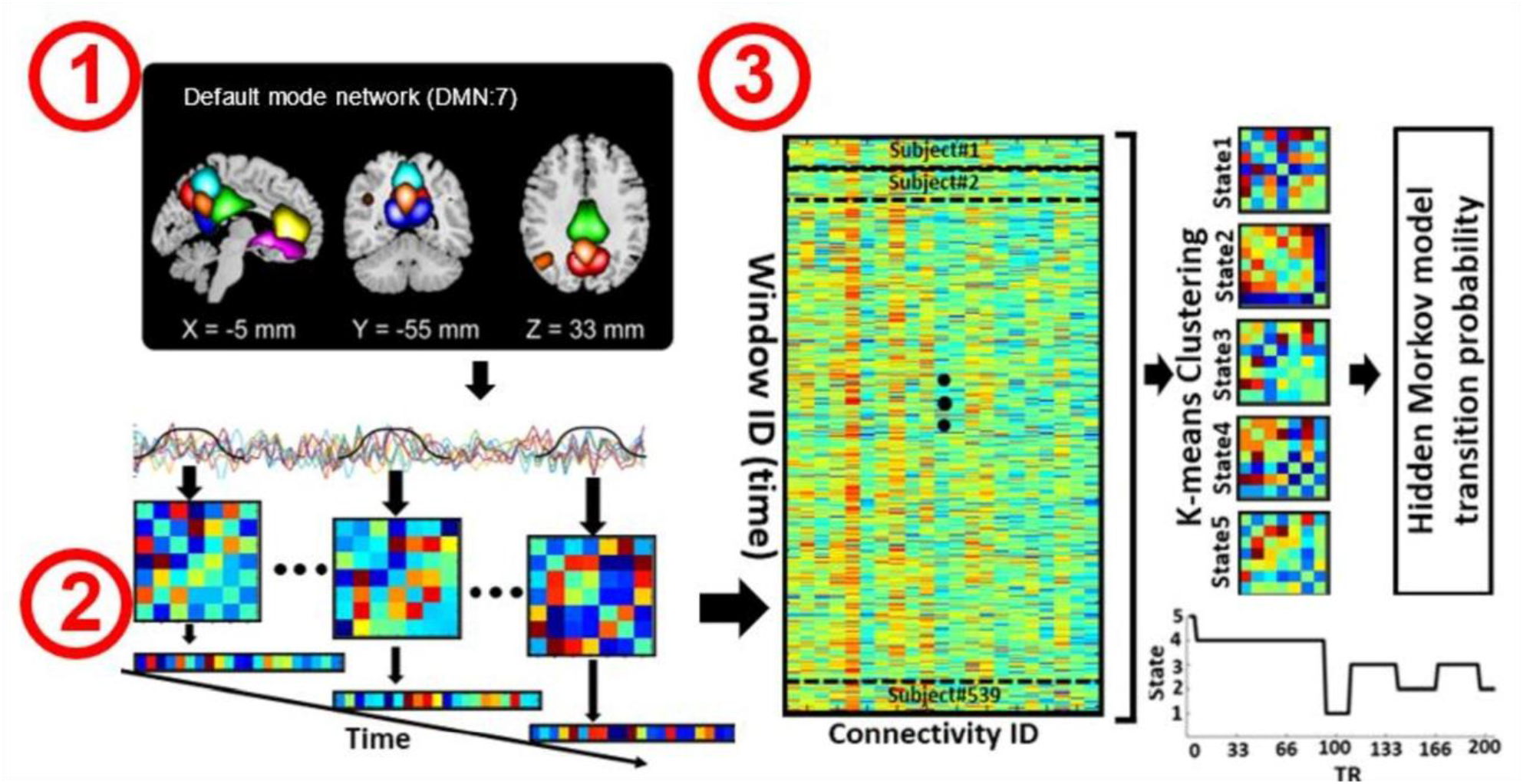
Analytic pipeline. Step1: The time-course signal of seven regions in the default mode network (DMN) has been identified using group-ICA. Step2: After identifying seven regions in the DMN, a taper sliding window was used to segment the time-course signals and calculate the FC matrix. Each FC matrix contains twenty-one connectivity features. Each feature represents the connectivity between a pair of DMN subnodes. Step3: After vectorizing the FC matrixes, we concatenated them applied and k-means clustering to group the FCs into five distinct clusters. Then, 25 hidden Markov model (HMM) features were calculated from the state vector of each subject. We investigated the association between HMM features and symptom severity in schizophrenia subjects.

### 2.4 Dynamic Functional Connectivity (dFC)

For each subject i = 1 … N, the dFC of the seven subnodes in the DMN was estimated via a sliding window approach, as shown in Figure 1. A tapered window obtained by convolving a rectangle (window size = 20 TRs or 40 s) with a Gaussian (σ = 3 s) was used to localize the dataset at each time point. A covariance matrix was calculated to measure the dFC (Figure 1 Step2). The dFC estimates of each window for each subject were concatenated to form a (C × C × T) array (where C = 7 denotes the number of subnodes, and T = 124 in COBRE and T = 137 in FBIRN denotes the number of windows), which represented the changes in brain connectivity between subnodes as a function of time (Allen et al., 2014; Calhoun et al., 2014; Fu et al., 2019). Since the temporal resolution and the eye condition of the two datasets were different, we did not combine them in our study and chose to analyze them separately instead.

### 2.5 Clustering and Latent Transition Feature Estimation

After calculating the dFC of each subject separately, we vectorized each FC window and concatenated all subjects, including both the SZ and HC groups, as shown in Step 3 of Figure 1. Next, the k-means clustering algorithm was applied to the dFC windows to partition the concatenated matrix into a set of distinct clusters or states (Allen et al., 2014; Calhoun et al., 2014; Zhi et al., 2018). An FC state, which is a conceptual analogy of an EEG microstate, is a global pattern of DMN connectivity that remains quasi-stable for a short period of time before changing to another connectivity pattern that also remains quasi-stable (Calhoun et al., 2014). The optimal number of centroid states was estimated to be 5 using the elbow criterion based on the ratio of within to between cluster distance. Correlation was implemented as a distance metric in the clustering algorithm in 1000 iterations. The output of k-means clustering includes five distinct states across all subjects and a state vector for each individual. The state vector shows how the DMN changes between each pair of states over time. Next, for each subject, we calculated the transition probability between states via an HMM, and this probability was used as a latent feature of the dFC. The transition probability, *a*_*ij*_, is the probability of the network to transition from state *j* at time *t* to state *i* at time *t+1*, (Step3 in Figure 1).

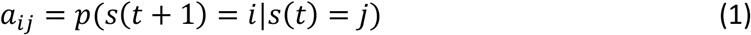

For each subject, twenty-five HMM features were obtained from the five states. This analysis was repeated separately for both the COBRE and FBIRN datasets, and the results of the analyses were compared to identify reoccurring patterns.

### 2.6 Quantifying Group Differences with a Feature Selection Method

Logistic regression (LR) classification was employed to quantify the difference between SZ subjects and HCs based on the twenty-one connectivity features of each state. The process is shown in Figure 2. In this process, the FC matrix of each window was converted to a vector. For the seven regions in the DMN, we obtained a total of twenty-one features (i.e., C_1_, C_2_, …, C_21_). Elastic net regularization (ENR), a machine learning-based feature selection method, was used to model the difference between the HC and SZ subjects (Zou and Hastie, 2005; Tibshirani, 2011). ENR applies both L1- and L2-regularization, as shown in Equation 2 and 3. In this method, the LR model parameters (i.e., feature coefficients) will move towards zero as *λ* increases. This will give a trajectory of the model parameters as a function of *λ* and form a model regularization path. The features related to the slowest decaying coefficients were considered to be most important. The cost function used in ENR is shown in the equations below:

**Figure 2.**
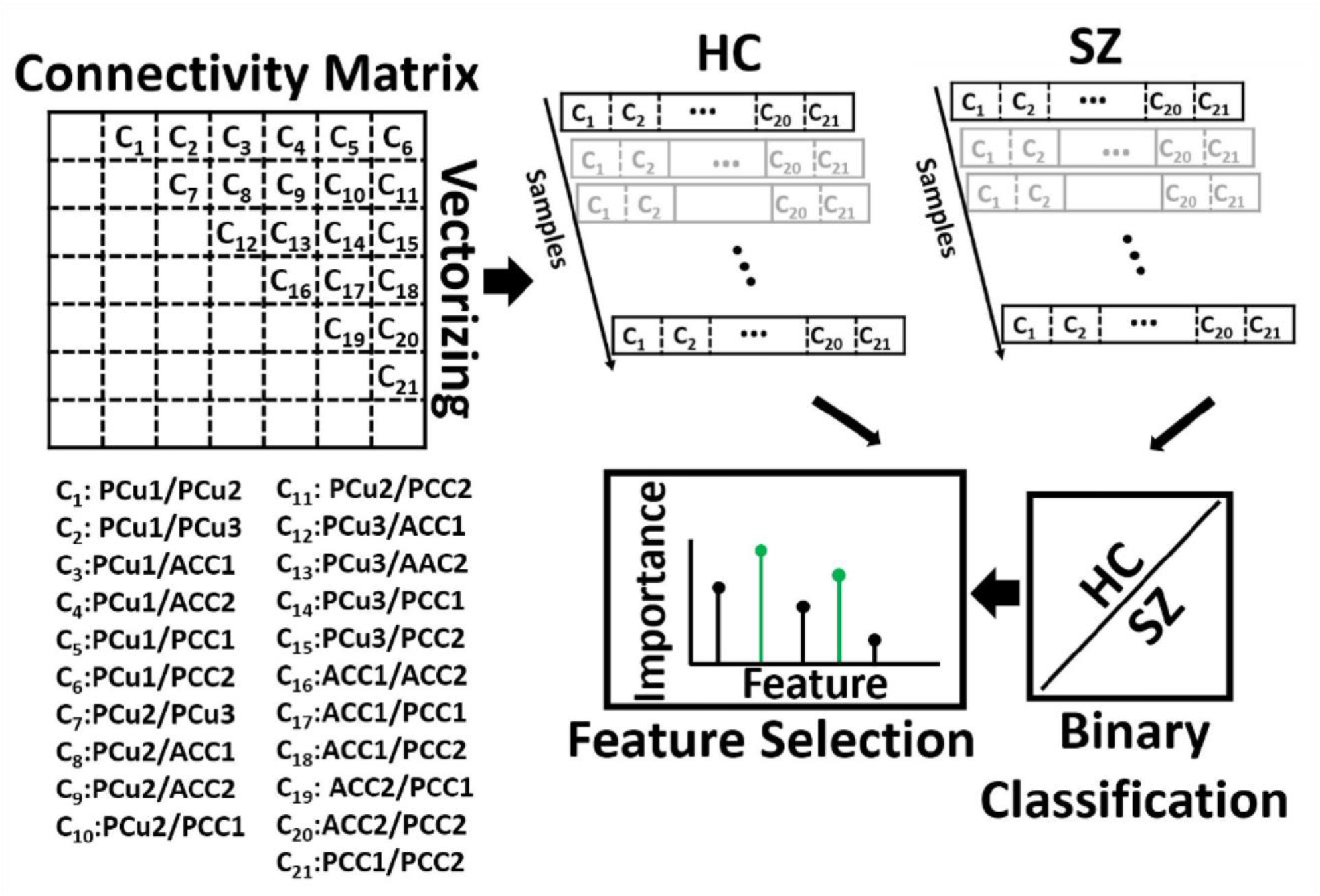
Feature selection. The connectivity features of seven default mode network (DMN) subnodes were used as inputs to fit a logistic regression classifier to discriminate SZs from HCs. With seven subnodes of the DMN, we had twenty-one connectivity features. The feature selection method, elastic net regularization (ENR), used the model generated by the classifier and the input features to identify the most predictive features. ACC: Anterior cingulate cortex, PCC: posterior cingulate cortex, PCu: Precuneus. Table 2 provides more information about different subnodes.

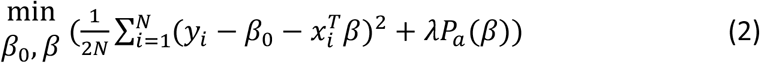

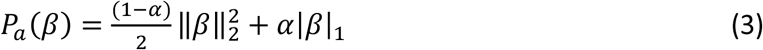

where *N* is the number of samples, *y*_*i*_ is the label of sample *i, x*_*i*_ is the feature vector of sample *i, β* and *β*_*0*_ are model parameters, *λ* is the regularization parameter, and P_a_(β) is the penalty term in which α (a scaler value) determines the relative contributions of the *L1* and *L2* norms where α = 1 is purely an *L1* norm and α = 0 is purely an *L2* norm (Zou and Hastie, 2005).

The LR model was fit using 10-fold nested cross-validation (CV) with a train-test ratio of 9:1 (Wainer and Cawley, 2018). In nested CV, the data was divided into training and test sets in an outer fold, while the training data was further divided into another training set and a validation set in the inner fold. The optimized parameters are obtained using the inner-loop training and validation data. Here, the hyperparameters of each model are tuned to minimize the inner-fold CV error of the generalization performance by sweeping the penalty parameter across 100 logarithmically-distributed values between 10^−5^ to 10^5^. Using the results of the performance of the classifiers upon the test data, we computed the receiver operating characteristic (ROC) of the cross-validation and the area under the curve (AUC) as a measure of separability between SZs and HCs. To identify the most informative feature to the classification between SZs and HCs, we calculated the proportion of models for which a given parameter was retained during the penalty parameter sweep in the inner fold. This measurement may be interpreted as the relative importance of each feature in the classification. We further applied multiple comparisons in a one-way analysis of variance (ANOVA) test and found the groups of features that most contributed to the model classifying between HC and SZ subjects.

### 2.7 Statistical Analysis

To find a link between the twenty-five HMM features and the PANSS of the SZ group, we used Pearson’s partial correlation method accounting for age (both datasets), gender (both datasets), and scanning site (for FBIRN only). All p values were adjusted by the Benjamini-Hochberg correction method for false discovery rate, or FDR (Yoav Benjamini; Yosef Hochberg, 1995).

## 3 Results

### 3.1 Dynamic functional connectivity states

Five states were identified in both datasets, as shown in Figure 3. For easier comparison, we vertically aligned the similar states of both datasets. The Pearson correlation between the cluster centroid matrix was used to quantify the similarity between the states identified within each dataset. The state centroid values are shown in Table 3. Similar dynamic DMN FC was observed in both datasets even though the eye condition during recording was different across datasets. The ACC regions showed negative connectivity in all states of both datasets except for state 5 in the FBIRN data. The connectivity between the ACC and PCC (ACC/PCC) was negative in all states of both datasets, and the connectivity between PCu and PCC was positive in all states of both datasets except state 3 of the FBIRN dataset. Within PCu, within PCC, and between PCu and ACC showed similar positive and negative connectivity patterns across datasets.

**Table 3.**
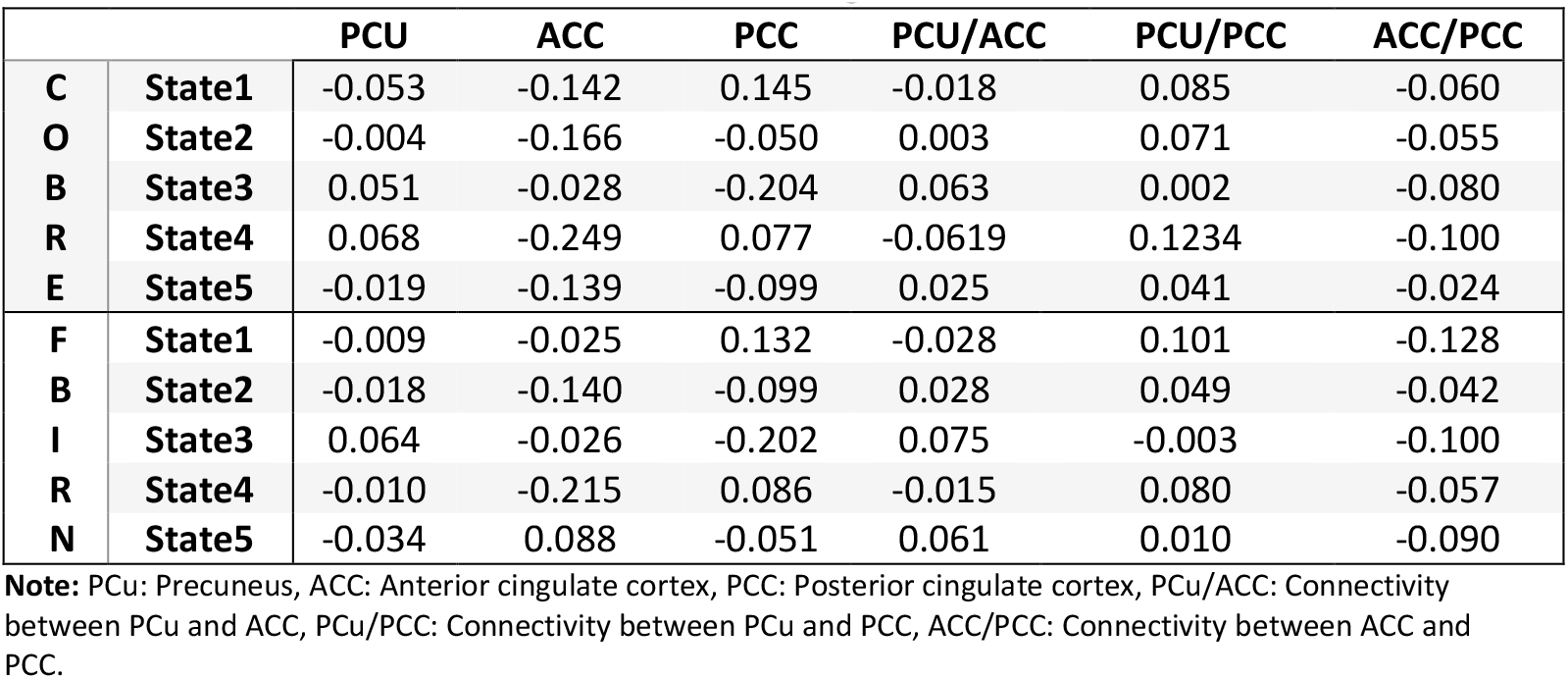
Mean value of the connectivity in each state based on the cluster centroid matrix from Figure 3

**Figure 3.**
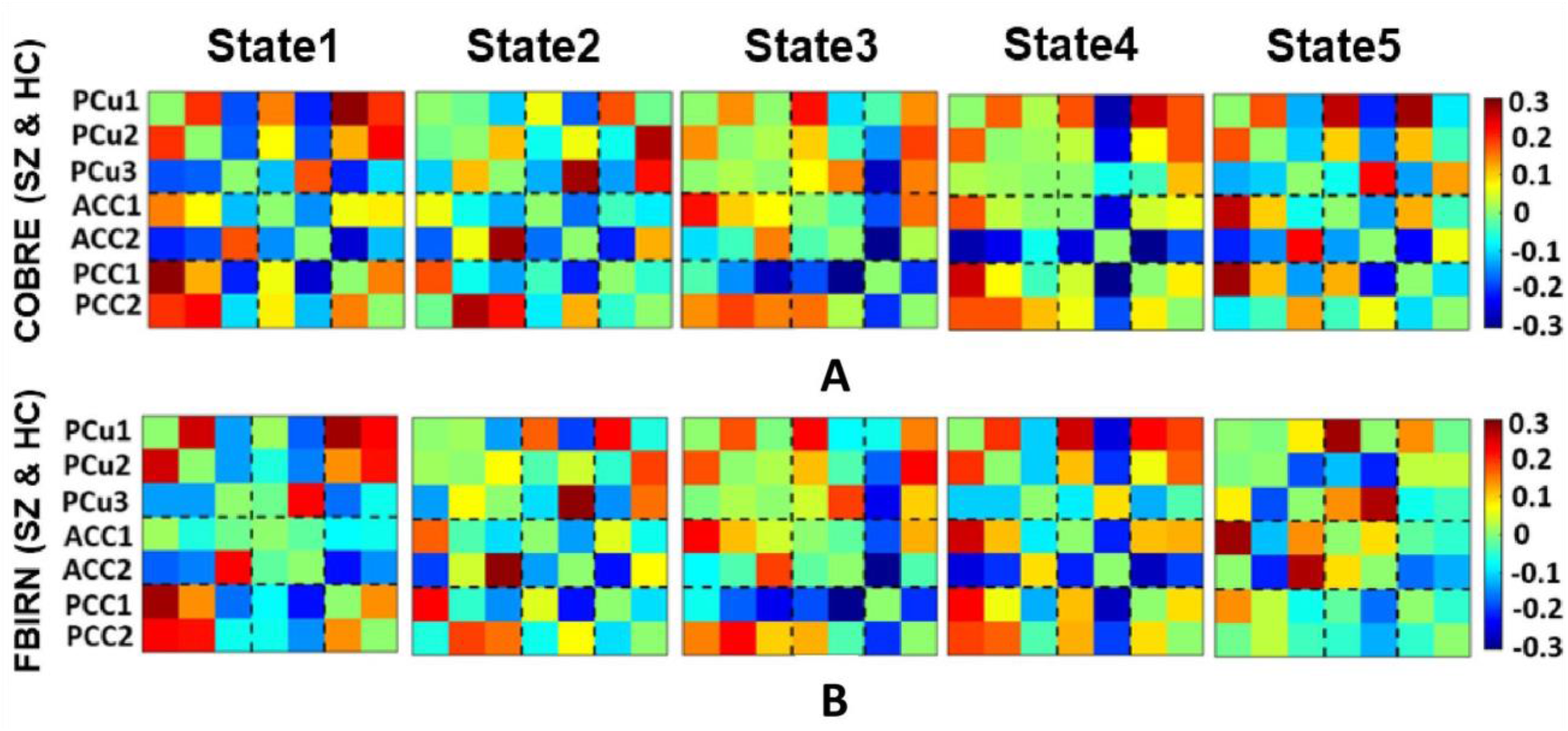
Dynamic connectivity states results. **A**) The five dFC states identified with k-means clustering in the COBRE data for both SZ and HC subjects. **B)** The five dFC states identified with k-means clustering in the FBIRN data for both SZ and HC subjects. The similar states between the two dataset are aligned vertically. The similarity between states was measured by the Pearson correlation of the cluster centroid matrix of the two datasets. There is not a similar pattern between COBRE and FBIRN in state 5. The colorbar shows the strength of the connectivity. Table 2 provides more information about different subnodes.

### 3.2 Difference between SZ and HC connectivity in each state

A feature learning method embedded in a 10-fold LR classifier was used to identify the differences between SZ and HC subjects in each state (Figure 2). Figures 4 and 5 show the classification and feature learning results of each state in the classification between SZ and HC subjects in the COBRE and FBIRN datasets, respectively. Multiple features were identified as equally important for differentiating each state. A detailed description of the feature learning results can be found in the section, “Classification and Feature Selection Results for Each State,” of the supplementary material. Figure 6 consolidates results from Figures 4 and 5 for easier comparison across datasets. It depicts differences in features between the SZ and HC groups (corrected p<0.05) that were selected by ENR in the COBRE dataset (Figure 6A) and the FBIRN dataset (Figure 6B). Red lines show stronger connectivity in HC subjects relative to SZ subjects, and blue lines show stronger connectivity in SZ subjects relative to HC subjects. The line width indicates the difference in connectivity strength between the SZ and HC groups.

**Figure 4.**
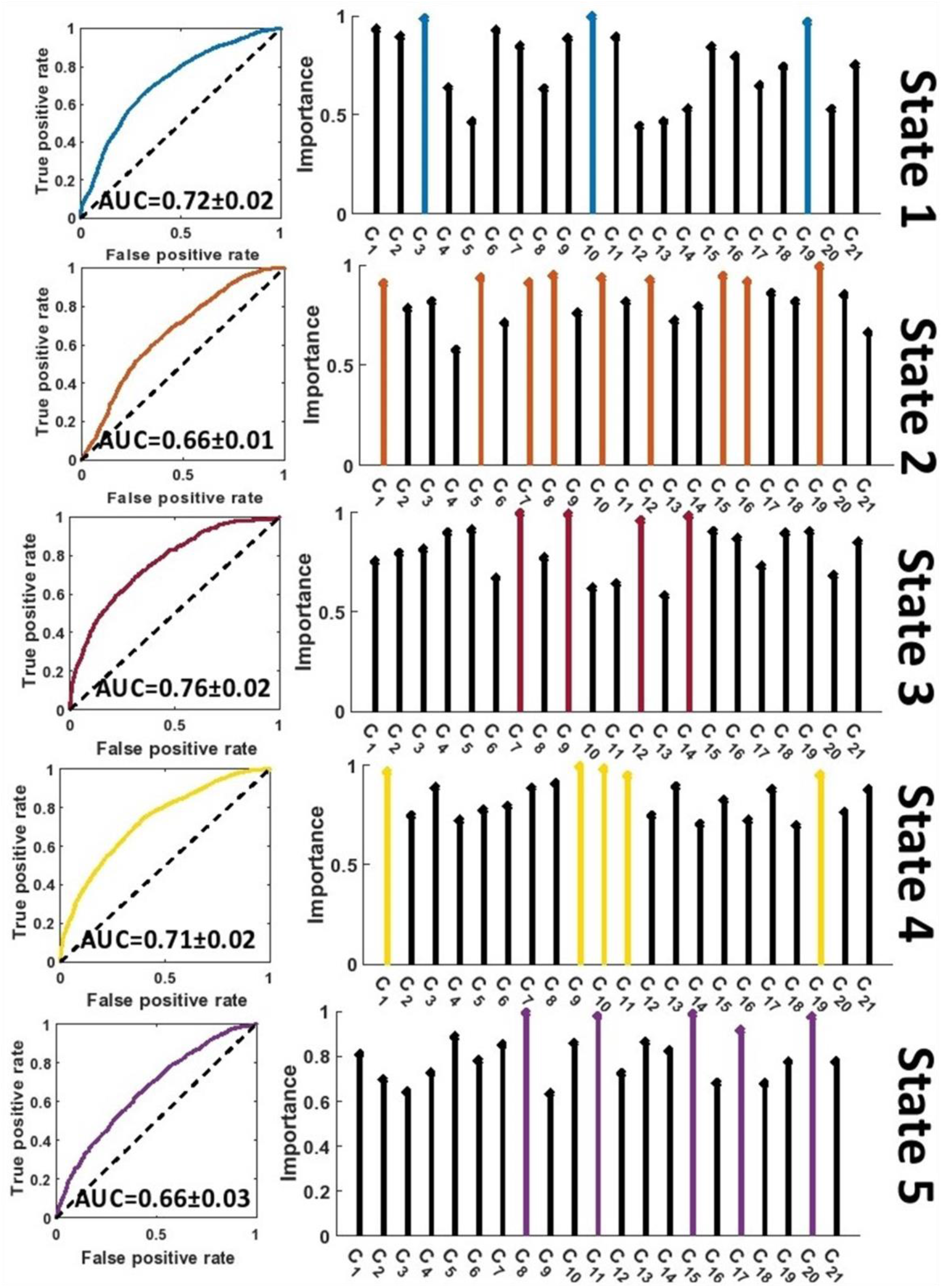
Feature selection results in COBRE dataset. The left panel shows the receiver operating characteristic curve of the classification between SZs and HCs in each state. The right panel shows the relative importance of the features to the classification. The colorful features are groups of equally important features that were found to be of greater importance than the remaining features by a multiple comparison ANOVA test. The features (C_1_ – C_21_) are defined in Figure 2. AUC: Area under the curve.

**Figure 5.**
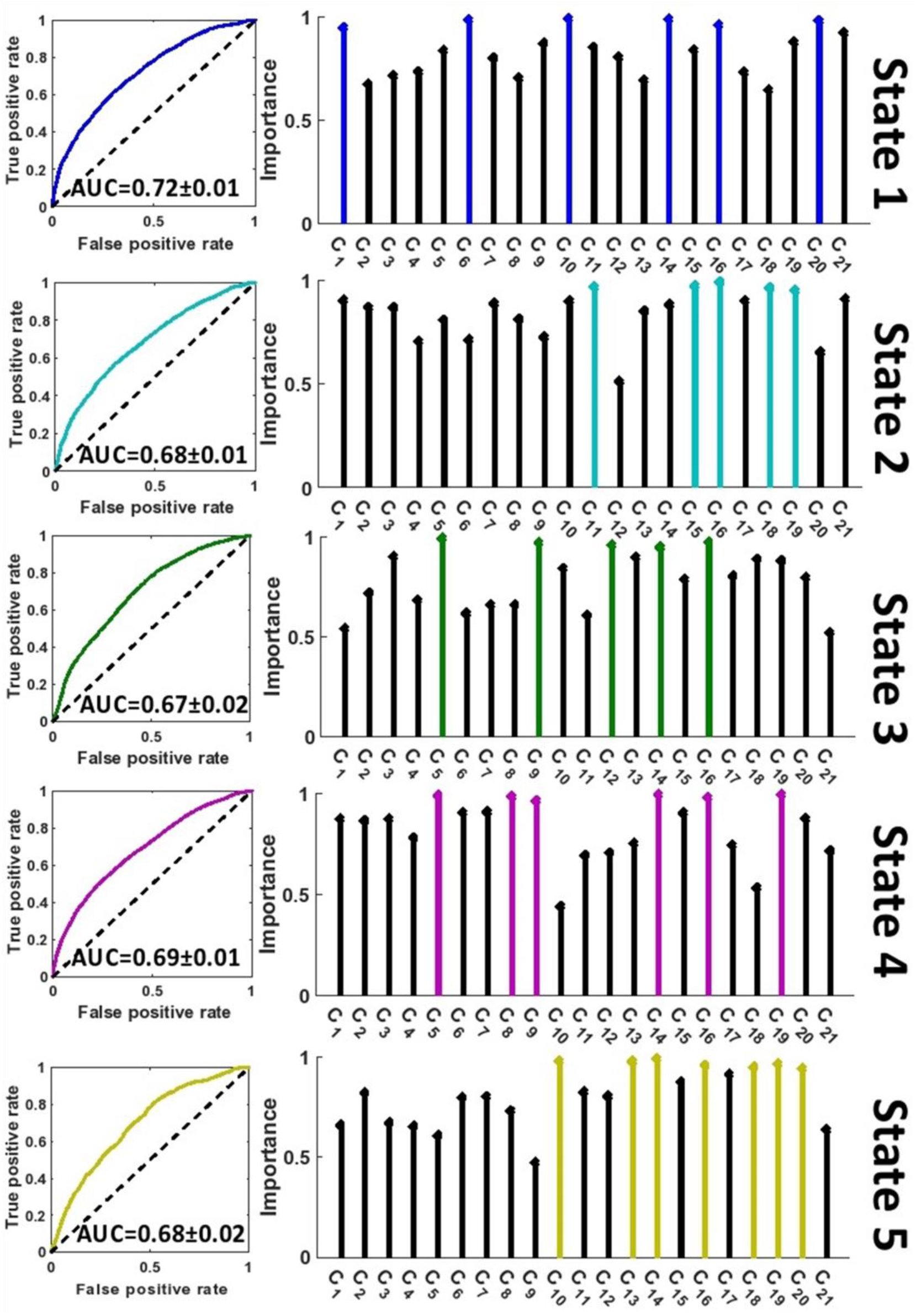
Feature selection results in FBIRN dataset. The left panel shows the receiver operating characteristic curve of the classification between SZs and HCs in each state. The right panel shows the relative importance of the features to the classification. The colorful features are groups of equally important features that were found to be of greater importance than the remaining features by a multiple comparison ANOVA test. The features (C_1_ – C_21_) are defined in Figure 2. AUC: Area under the curve.

**Figure 6.**
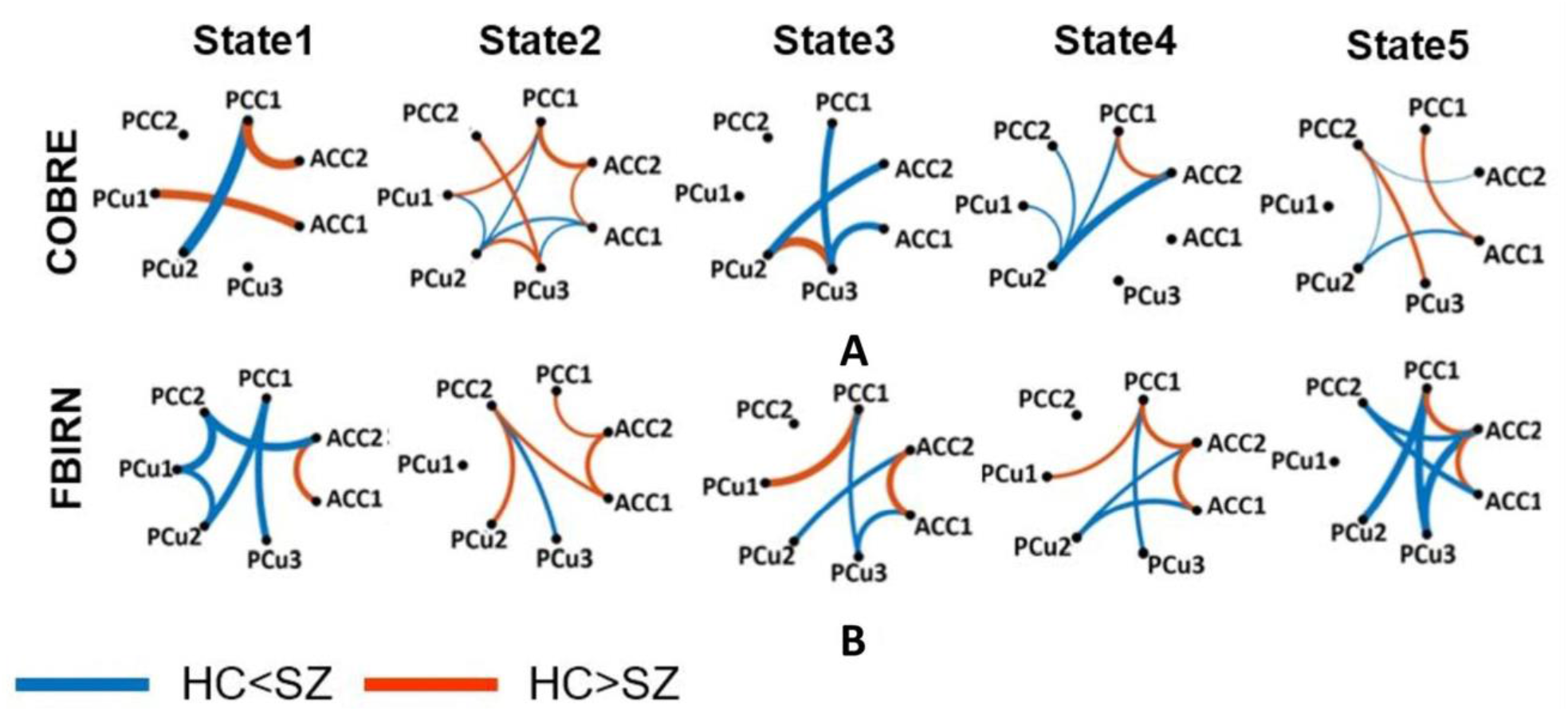
Group difference between SZ and HC connectivity in each state. Group differences in dFC of those connectivity features selected by elastic net regularization method (see Figures 4 and 5) in each state (corrected p < 0.05). Wider line means larger group difference. Red lines represent increased connectivity while blue lines represent decreased connectivity in HC subjects. a COBRE dataset. b FBIRN dataset. ACC: Anterior cingulate cortex, PCC: Posterior cingulate cortex, PCu: Precuneus. HC: Healthy control, SZ: Schizophrenia. Table 2 provides more information about different subnodes.

Disrupted connectivity between the PCu and PCC (PCu/PCC) was observed in both datasets. In both datasets, we observed higher PCu/PCC connectivity in SZ subjects in states 1 and 4 (corrected p<0.05). State 5 of the FBIRN dataset also displayed higher PCu/PCC in SZ subjects. In the COBRE dataset, SZ subjects showed a lower PCu/PCC connectivity in state 2 and state 5 (corrected p<0.05), and in the FBIRN dataset, PCu/PCC connectivity of SZ was lower in state 3 (corrected p<0.05). Also, for both datasets, the connectivity between PCu and the cingulate cortex (including both the ACC and PCC) of SZ subjects was higher in state 1, state 3, state 4, and state 5 (corrected p<0.05).

Both datasets showed higher ACC connectivity in HC subjects in state 2 (corrected p<0.05), and the FBIRN data showed a higher ACC connectivity in HC subjects in states 1, 3, 4, and 5 (corrected p<0.05). Higher HC PCC/ACC connectivity was observed in states 2 and 4 of both datasets (corrected p<0.05), and higher HC PCC/ACC connectivity was also observed in states 1 and 5 of the COBRE dataset (corrected p<0.05). For state 3 in both datasets, PCC/ACC connectivity was similar across HC and SZ groups. Additionally, PCC connectivity of the HC and SZ groups was similar across all states and both datasets.

### 3.3 Symptom correlation with HMM features

It is important to understand how the dynamic aspects of DMN connectivity correlate with symptom severity. In the COBRE dataset, one correlation between total PANSS and an HMM feature was significant after accounting for FDR correction (FDR corrected p < 0.05). In this instance, symptom severity showed a positive correlation with transitions from state 4 to state 2 (r=0.40, FDR corrected p=0.02, n=64). Similar results were found in the FBIRN data in which the transition probability from state 4 to state 2 showed a significant correlation with negative PANSS (r=0.32, FDR corrected p=0.002, n=141).

## 4 Discussion

Two key goals of the current study were (1) ensuring the generalizability of results by identifying similar patterns found in two distinct datasets and (2) offering an explanation for preexisting contradictory findings on DMN connectivity in schizophrenia. We explored the temporal dynamics of functional connectivity among several data-driven DMN subnodes from the PCC, ACC, and PCu regions using rs-fMRI of two schizophrenia datasets. We further explored SZ and HC group connectivity differences among the subnodes, identifying multiple patterns that generalized across datasets.

In both datasets, we observed negative connectivity within the ACC (except state 5 of FBIRN) and between the ACC and PCC of all states of both datasets. While the connectivity between the PCu and PCC was positive in all states of both datasets except state 3 of the FBIRN dataset. On the other hand, the connectivity between the PCu and ACC, within the PCu, and within the PCC demonstrated a similar pattern in both datasets, fluctuating between positive and negative connectivity. Here, using data-driven subnodes within the DMN, we showed that the brain network is highly dynamic. Previous literature typically ignored this dynamical DMN behavior. In contrast to the previous study that evaluated DMN dynamics using pre-defined regions of interest (Du et al., 2016), the work presented here is the first study that utilized data-driven subnodes, compared the within-DMN connectivity between SZ and HC subjects, and linked the temporal patterns of the DMN with symptom severity in SZ subjects. As recent work has emphasized, it is essential to ensure that data within a node is consistent; otherwise, the results can be misleading (Yu et al., 2017). This is especially true when studying dynamics (Iraji et al., 2020). The Neuromark pipeline that we used to identify subnodes yields reproducible nodes that should contribute to the overall generalizability of our results (Du et al., 2020).

Previously, a few studies directly examined ACC functional connectivity in the pathophysiology of schizophrenia. However, inconsistent results were observed. One study reported lower ACC connectivity in SZ (number of subject or N=58) subjects relative to HCs (N=61) (Shukla et al., 2019). A recent study showed a higher ACC connectivity for SZ (N=32) subjects at baseline relative to HCs (N=32) and a decreased ACC connectivity after one week of olanzapine treatment (Li et al., 2019). In the current study, we identified a pattern of disrupted ACC connectivity in the smaller dataset (i.e., COBRE), in which one state showed a higher ACC connectivity in HCs and other states showed no significant differences between HC and SZ groups. On the other hand, in the FBIRN dataset, which is a relatively large dataset compared to the COBRE dataset and the datasets in the studies mentioned above, we found a consistent increased ACC connectivity of HC subjects in all states. A possible explanation of previous inconsistent findings is the small sample size of the studies. However, even in the smaller dataset, we highlighted increased ACC connectivity in HCs with the dFC approach. As such, the use of sFC obtained from unconstrained rs-fMRI could be another explanation for previous inconsistent results on ACC connectivity. Finally, a previous study in a relatively small number of subjects (N=13) reported marginally (p=0.05) greater within-PCC connectivity in SZs relative to HCs (Whitfield-Gabrieli et al., 2009). However, in the current study, in which both datasets had a relatively larger sample size, no significant differences within-PCC were observed in any state. This supports the importance of using data-driven subnodes to study within-PCC connectivity in schizophrenia pathophysiology.

Although most previous studies of DMN functional connectivity focused on the ACC and PCC, we further highlighted the role of PCu/PCC connectivity via a comparison between HC and SZ subjects in two different datasets. In three of five states of both datasets, we found that the PCu/PCC connectivity was greater in SZs than HCs. However, we observed unique behavior across dFC states that would not be captured by sFC. Using sFC, previous studies reported both increases (Whitfield-Gabrieli et al., 2009; Peeters et al., 2015) and decreases (Wang et al., 2014) in the PCu/PCC connectivity in schizophrenia. These contradictory results are possibly due to focusing on sFC and averaging the functional connectivity across time. The current study showed a disrupted pattern of PCu/PCC connectivity with a relatively large sample and potentially highlighted the importance of studying functional connectivity sampled from shorter periods.

We investigated the link between symptom severity and dFC temporal patterns in each subject. Consistent across both datasets, we found a significant positive correlation between symptom severity and the transition from a state with low PCu/PCC and high ACC connectivity to a state with higher PCu/PCC and lower ACC connectivity. These results emphasize the role of cingulate cortex connectivity and PCu/PCC connectivity as potential biomarkers of SZ, and the role is further highlighted in the more severe SZ subjects. A previous study explored the link between dFC features such as the number of transitions between states and the dwell times of each state, and the results were not significant after FDR correction (Rabany et al., 2019). Our current study shows that HMM features extracted from dFC are correlated with symptom severity and supports the importance of exploiting the network dynamics as potential biomarkers. This also motivates future work studying the relationship of symptom severity to other dFC features.

The current study extends previous studies performed on the same datasets that investigated the dynamics of the whole-brain network connectivity (Damaraju et al., 2014; Sendi et al., 2020). In a larger brain network, a group of brain networks such as the visual, sensorimotor, and auditory networks, which are strongly correlated, may mask less-correlated networks and limit spatiotemporal resolution (Schlesinger et al., 2017). That could potentially delineate why the main results of these studies focused on these dominant networks and reported less on networks like the DMN that may have been masked. Also, due to higher DMN activity during resting state, studying the dynamics of this network can reveal new information that cannot be found by analyzing the whole-brain connectivity. Although in the current study we focused on the DMN because of prior knowledge of the role of the network in the pathophysiology of schizophrenia, future investigations and methods that can mechanistically remove irrelevant networks are needed (Cohen et al., 2015; Schlesinger et al., 2017; Qiao et al., 2019).

Finally, as mentioned earlier, the eye condition is different in the COBRE and FBIRN datasets. A previous study reported that different eye conditions might modulate DMN dynamics (Zhang et al., 2018), which could explain some differences in the DMN dynamics between the two datasets. State 5 of FBIRN dataset was distinguished from all other states in both datasets by showing higher within-ACC connectivity. Since previous literature showed higher activity in the ACC during sleep (Hobson and Pace-Schott, 2002), we wonder whether this connectivity pattern is possibly linked to the light sleep or drowsiness that may have occurred during the unconstrained state of eyes-closed in the FBIRN dataset. This potentially demonstrated another benefit of dynamic functional connectivity analysis, separating undesired states from the rest, specifically when the eye is closed.

### 4.1 Limitations

There are some limitations to this work. Symptom scores are highly dependent on the skill and knowledge of the rater and the inclination of the subjects to be accurate in describing their symptoms (Kay et al., 1987). As such, our use of the FBIRN dataset, which was collected from multiple sites and raters, may have introduced a degree of bias into our analyses. The choice of window size is an implicit assumption about the dynamic behavior of the network in that a short window captures more rapid fluctuations, whereas a longer window causes more smoothing. Previous studies suggest that a window size between 30-60 s provides a reasonable choice for capturing dFC variation (Preti et al., 2017). The duration of scanning was over 5 minutes, which has been shown to result in reliable and replicable resting-state FNC (Van Dijk et al., 2010; Abrol et al., 2017). While we are encouraged by the similarity of results across multiple data sets, schizophrenia is likely a heterogeneous disorder, and more work is needed to evaluate the potential of multiple types of connectivity patterns within this group to provide additional insight into the disorder.

### 4.2 Conclusion

Previous studies focused on static connectivity of the DMN, including the PCC, ACC, and PCu and showed an essential role of this connectivity in schizophrenia. In the current work, we extended this existing body of research into the domain of dynamics by investigating the temporal patterns of connectivity in the DMN. A comparison of the DMN connectivity in SZs and HCs identified patterns of disruption in a shorter timescale that were reproducible across two relatively large datasets with distinct collection protocols. These patterns of disruption could possibly explain why previous studies of DMN connectivity showed contradictory results. In both datasets, we found that SZ subjects with higher symptom severity are more likely to transition from a state with lower PCu/PCC connectivity and higher ACC connectivity to a state with higher PCu/PCC connectivity and lower ACC connectivity. This highlights the potential relationship between symptom severity and the dysregulation of the dynamical properties of DMN functional connectivity.

## 5 Data Availability Statement

The datasets supporting the findings of the study are available upon a reasonable request made to the corresponding author. The code used for preprocessing and dFNC calculation are available at https://trendscenter.org/software/.

## 6 Conflict of Interest

Dr. Mathalon is a consultant for Boehringer Ingelheim, Cadent Therapeutics, and Greenwich Biosciences.

## 7 Author Contributions

Mohammad Sendi and Elaheh Zendehrouh analyzed the data. Zening Fu preprocessed the imaging data. Mohammad Sendi, Charles Ellis, Jessica Turner, and Vince Calhoun wrote the manuscript. Daniel Mathalon, Judith Ford, Adrian Preda, Theo van Erp, Robyn Miller, and Godfrey Pearlson provided constructive feedback on the manuscript during its preparation. All coauthors reviewed the manuscript before submission.

## 8 Funding

This work is funded by NIH grants: R01MH094524, R01MH119069, R01MH118695, R01EB020407, and R01MH121101 and by NSF grant IIS-1631819.

## 9 Acknowledgments

We thank the study participants who made our findings possible.

## 10 Ethics Statement

Data were obtained from the Mind Research Network Center of Biomedical Research Excellence (COBRE) and the FBIRN projects. The FBIRN raw imaging data were collected from seven sites including the University of California, Irvine; the University of California, Los Angeles; the University of California, San Francisco; Duke University/the University of North Carolina at Chapel Hill; the University of New Mexico; the University of Iowa; and the University of Minnesota. In this study, written informed consent was obtained from all participants. Institutional review boards approved the consent process of each study site.

## Reference

Abrol, A., Damaraju, E., Miller, R. L., Stephen, J. M., Claus, E. D., Mayer, A. R., et al. (2017). Replicability of time-varying connectivity patterns in large resting state fMRI samples. Neuroimage 163, 160–176. doi:10.1016/j.neuroimage.2017.09.020.

Aine, C. J., Bockholt, H. J., Bustillo, J. R., Cañive, J. M., Caprihan, A., Gasparovic, C., et al. (2017). Multimodal Neuroimaging in Schizophrenia: Description and Dissemination. Neuroinformatics 15, 343–364. doi:10.1007/s12021-017-9338-9.

Allen, E. A., Damaraju, E., Plis, S. M., Erhardt, E. B., Eichele, T., and Calhoun, V. D. (2014). Tracking whole-brain connectivity dynamics in the resting state. Cereb. Cortex 24, 663–676. doi:10.1093/cercor/bhs352.

Bhinge, S., Long, Q., Calhoun, V. D., and Adali, T. (2019). Spatial Dynamic Functional Connectivity Analysis Identifies Distinctive Biomarkers in Schizophrenia. Front. Neurosci. 13. doi:10.3389/fnins.2019.01006.

Calabrese, D. R., Wang, L., Harms, M. P., Ratnanather, J. T., Barch, D. M., Cloninger, C. R., et al. (2008). Cingulate gyrus neuroanatomy in schizophrenia subjects and their non-psychotic siblings. Schizophr. Res. 104, 61–70. doi:10.1016/j.schres.2008.06.014.

Calhoun, V. D., Miller, R., Pearlson, G., and Adali, T. (2014). The Chronnectome: Time-Varying Connectivity Networks as the Next Frontier in fMRI Data Discovery. Neuron 84, 262–274. doi:10.1016/j.neuron.2014.10.015.

Chain, M., Carlo, M., and Carlo, M. (2019). Individual Variation in Brain Network Topography is Linked to Schizophrenia Symptomatology. https://doi.org/10.1101/692186, 1–31. doi:10.1101/541979.

Cohen, M. B., Elder, S., Musco, C., Musco, C., and Persu, M. (2015). Dimensionality reduction for kmeans clustering and low rank approximation. Arxiv Prepr. arXiv1410.6801. doi:10.1145/2746539.2746569.

Damaraju, E., Allen, E. A., Belger, A., Ford, J. M., McEwen, S., Mathalon, D. H., et al. (2014). Dynamic functional connectivity analysis reveals transient states of dysconnectivity in schizophrenia. NeuroImage Clin. 5, 298–308. doi:10.1016/j.nicl.2014.07.003.

Du, Y., Fu, Z., Sui, J., Gao, S., Xing, Y., Lin, D., et al. (2019). NeuroMark: an adaptive independent component analysis framework for estimating reproducible and comparable fMRI biomarkers among brain disorders. medRxiv, 19008631. doi:10.1101/19008631.

Du, Y., Fu, Z., Sui, J., Gao, S., Xing, Y., Lin, D., et al. (2020). NeuroMark: An automated and adaptive ICA based pipeline to identify reproducible fMRI markers of brain disorders. NeuroImage Clin. 28, 102375. doi:10.1016/j.nicl.2020.102375.

Du, Y., He, H., Wu, L., Yu, Q., Sui, J., and Calhoun, V. D. (2015). Dynamic default mode network connectivity diminished in patients with schizophrenia. Proc. - Int. Symp. Biomed. Imaging 2015-July, 474–477. doi:10.1109/ISBI.2015.7163914.

Du, Y., Pearlson, G. D., Yu, Q., He, H., Lin, D., Sui, J., et al. (2016). Interaction among subsystems within default mode network diminished in schizophrenia patients: A dynamic connectivity approach. Schizophr. Res. 170, 55–65. doi:10.1016/j.schres.2015.11.021.

Engels, G., Vlaar, A., McCoy, B., Scherder, E., and Douw, L. (2018). Dynamic Functional Connectivity and Symptoms of Parkinson’s Disease: A Resting-State fMRI Study. Front. Aging Neurosci. 10, 1–9. doi:10.3389/fnagi.2018.00388.

First, M. B., Spitzer, R. L., Gibbon, M., and Williams, J. B. W. (2002a). Structured Clinical Interview for DSM-IV-TR Axis I Disorders, Research Version, Non-patient Edition. (SCID-I/NP).

First, M. B., Spitzer, R. L., Gibbon, M., and Williams, J. B. W. (2002b). Structured Clinical Interview for DSM-IV-TR Axis I Disorders, Research Version, Patient Edition. (SCID-I/P). New York Biometrics Res.

Fu, Z., Tu, Y., Di, X., Du, Y., Sui, J., Biswal, B. B., et al. (2019). Transient increased thalamic-sensory connectivity and decreased whole-brain dynamism in autism. Neuroimage 190, 191–204. doi:10.1016/j.neuroimage.2018.06.003.

Garrity, A. G., Pearlson, G. D., MacKiernan, K., Lioyd, D., Kiehl, K. A., and Calhoun, V. D. (2007). Aberrant “Default Mode” Functional Connectivity in Schizophrenia. Am J Psychiatry 164, 450–457.

Guo, W., Liu, F., Chen, J., Wu, R., Li, L., Zhang, Z., et al. (2017). Hyperactivity of the default-mode network in first-episode, drug-naive schizophrenia at rest revealed by family-based case-control and traditional case-control designs. Med. (United States) 96. doi:10.1097/MD.0000000000006223.

Hare, S. M., Ford, J. M., Ahmadi, A., Damaraju, E., Belger, A., Bustillo, J., et al. (2017). Modality-Dependent Impact of Hallucinations on Low-Frequency Fluctuations in Schizophrenia. Schizophr. Bull. 43, 389–396. doi:10.1093/schbul/sbw093.

Hare, S. M., Ford, J. M., Mathalon, D. H., Damaraju, E., Bustillo, J., Belger, A., et al. (2019). Salience-default mode functional network connectivity linked to positive and negative symptoms of schizophrenia. Schizophr. Bull. 45, 892–901. doi:10.1093/schbul/sby112.

Hobson, J. A., and Pace-Schott, E. F. (2002). The cognitive neuroscience of sleep: Neuronal systems, consciousness and learning. Nat. Rev. Neurosci. 3, 679–693. doi:10.1038/nrn915.

Hu, M. L., Zong, X. F., Mann, J. J., Zheng, J. J., Liao, Y. H., Li, Z. C., et al. (2017). A Review of the Functional and Anatomical Default Mode Network in Schizophrenia. Neurosci. Bull. 33, 73–84. doi:10.1007/s12264-016-0090-1.

Iraji, A., Miller, R., Adali, T., and Calhoun, V. D. (2020). Space: A Missing Piece of the Dynamic Puzzle. Trends Cogn. Sci. 24, 135–149. doi:10.1016/j.tics.2019.12.004.

Kay, S. R., Fiszbein, A., and Opler, L. A. (1987). The positive and negative syndrome scale (PANSS) for schizophrenia. Schizophr. Bull. 13, 261–276. doi:10.1093/schbul/13.2.261.

Leech, R., and Sharp, D. J. (2014). The role of the posterior cingulate cortex in cognition and disease. Brain 137, 12–32. doi:10.1093/brain/awt162.

Li, H., Ou, Y., Liu, F., Chen, J., Zhao, J., Guo, W., et al. (2019). Reduced connectivity in anterior cingulate cortex as an early predictor for treatment response in drug-naive, first-episode schizophrenia: A global-brain functional connectivity analysis. Schizophr. Res., 1–7. doi:10.1016/j.schres.2019.09.003.

Liemburg, E. J., van der Meer, L., Swart, M., Curcic-Blake, B., Bruggeman, R., Knegtering, H., et al. (2012). Reduced connectivity in the self-processing network of schizophrenia patients with poor insight. PLoS One 7, 1–9. doi:10.1371/journal.pone.0042707.

Lynall, M. E., Bassett, D. S., Kerwin, R., McKenna, P. J., Kitzbichler, M., Muller, U., et al. (2010). Functional connectivity and brain networks in schizophrenia. J. Neurosci. 30, 9477–9487. doi:10.1523/JNEUROSCI.0333-10.2010.

Peeters, S. C. T., Van De Ven, V., Gronenschild, E. H. B. M., Patel, A. X., Habets, P., Goebel, R., et al. (2015). Default mode network connectivity as a function of familial and environmental risk for psychotic disorder. PLoS One 10, 1–19. doi:10.1371/journal.pone.0120030.

Preti, M. G., Bolton, T. A., and Van De Ville, D. (2017). The dynamic functional connectome: State-of-the-art and perspectives. Neuroimage 160, 41–54. doi:10.1016/j.neuroimage.2016.12.061.

Qiao, C., Gao, B., Lu, L. J., Calhoun, V. D., and Wang, Y. P. (2019). Two-step feature selection for identifying developmental differences in resting fMRI intrinsic connectivity networks. Appl. Sci. 9, 1–18. doi:10.3390/app9204298.

Rabany, L., Brocke, S., Calhoun, V. D., Pittman, B., Corbera, S., Wexler, B. E., et al. (2019). Dynamic functional connectivity in schizophrenia and autism spectrum disorder: Convergence, divergence and classification. NeuroImage Clin. 24, 101966. doi:10.1016/j.nicl.2019.101966.

Rashid, B., Arbabshirani, M. R., Damaraju, E., Cetin, M. S., Miller, R., Pearlson, G. D., et al. (2016). Classification of schizophrenia and bipolar patients using static and dynamic resting-state fMRI brain connectivity. Neuroimage 134, 645–657. doi:10.1016/j.neuroimage.2016.04.051.

Sanfratello, L., Houck, J. M., and Calhoun, V. D. (2019). Dynamic Functional Network Connectivity in Schizophrenia with Magnetoencephalography and Functional Magnetic Resonance Imaging: Do Different Timescales Tell a Different Story? Brain Connect. 9, 251–262. doi:10.1089/brain.2018.0608.

Schlesinger, K. J., Turner, B. O., Grafton, S. T., Miller, M. B., and Carlson, J. M. (2017). Improving resolution of dynamic communities in human brain networks through targeted node removal. PLoS One 12, 1–28. doi:10.1371/journal.pone.0187715.

Schumacher, J., Peraza, L. R., Firbank, M., Thomas, A. J., Kaiser, M., Gallagher, P., et al. (2019). Dynamic functional connectivity changes in dementia with Lewy bodies and Alzheimer’s disease. NeuroImage Clin. 22, 101812. doi:10.1016/j.nicl.2019.101812.

Sendi, M. S. E., Member, S., Zendehrouh, E., Fu, Z., Mahmoudi, B., Miller, R. L., et al. (2020). A Machine Learning Model for Exploring Aberrant Functional Network Connectivity Transition in Schizophrenia. in IEEE Southwest Symposium on IEEE Southwest Symposium on, 112–115.

Shukla, D. K., Wijtenburg, S. A., Chen, H., Chiappelli, J. J., Kochunov, P., Hong, L. E., et al. (2019). Anterior cingulate glutamate and GABA associations on functional connectivity in schizophrenia. Schizophr. Bull. 45, 647–658. doi:10.1093/schbul/sby075.

Skåtun, K. C., Kaufmann, T., Doan, N. T., Alnæs, D., Córdova-Palomera, A., Jönsson, E. G., et al. (2017). Consistent Functional Connectivity Alterations in Schizophrenia Spectrum Disorder: A Multisite Study. Schizophr. Bull. 43, 914–924. doi:10.1093/schbul/sbw145.

Stevens, F. L., Hurley, R. A., and Taber, K. H. (2011). Anterior cingulate cortex: Unique role in cognition and emotion. J. Neuropsychiatry Clin. Neurosci. 23, 121–125. doi:10.1176/jnp.23.2.jnp121.

Tibshirani, R. (2011). Regression shrinkage and selection via the lasso?: a retrospective. J. R. Stat. Soc. Ser. B Stat. Methodol. 73, 273–282.

Tzourio-Mazoyer, N., Landeau, B., Papathanassiou, D., Crivello, F., Etard, O., Delcroix, N., et al. (2002). Automated anatomical labeling of activations in SPM using a macroscopic anatomical parcellation of the MNI MRI single-subject brain. Neuroimage 15, 273–289. doi:10.1006/nimg.2001.0978.

Van Dijk, K. R. A., Hedden, T., Venkataraman, A., Evans, K. C., Lazar, S. W., and Buckner, R. L. (2010). Intrinsic functional connectivity as a tool for human connectomics: Theory, properties, and optimization. J. Neurophysiol. 103, 297–321. doi:10.1152/jn.00783.2009.

van Erp, T. G. M., Preda, A., Turner, J. A., Callahan, S., Calhoun, V. D., Bustillo, J. R., et al. (2015). Neuropsychological profile in adult schizophrenia measured with the CMINDS. Psychiatry Res. 230, 826–834. doi:10.1016/j.psychres.2015.10.028.

Vergara, V. M., Mayer, A. R., Kiehl, K. A., and Calhoun, V. D. (2018). Dynamic functional network connectivity discriminates mild traumatic brain injury through machine learning. NeuroImage Clin. 19, 30–37. doi:10.1016/j.nicl.2018.03.017.

Wainer, J., and Cawley, G. (2018). Nested cross-validation when selecting classifiers is overzealous for most practical applications. 1–9. Available at: http://arxiv.org/abs/1809.09446.

Wang, D., Zhou, Y., Zhuo, C., Qin, W., Zhu, J., Liu, H., et al. (2015). Altered functional connectivity of the cingulate subregions in schizophrenia. Transl. Psychiatry 5. doi:10.1038/tp.2015.69.

Wang, L., Hosakere, M., Trein, J. C. L., Miller, A., Ratnanather, J. T., Barch, D. M., et al. (2007). Abnormalities of cingulate gyrus neuroanatomy in schizophrenia. Schizophr. Res. 93, 66–78. doi:10.1016/j.schres.2007.02.021.

Wang, X., Xia, M., Lai, Y., Dai, Z., Cao, Q., Cheng, Z., et al. (2014). Disrupted resting-state functional connectivity in minimally treated chronic schizophrenia. Schizophr. Res. 156, 150–156. doi:10.1016/j.schres.2014.03.033.

Wang, Y., Tang, W., Fan, X., Zhang, J., Geng, D., Jiang, K., et al. (2017). Resting-state functional connectivity changes within the default mode network and the salience network after antipsychotic treatment in early-phase schizophrenia. Neuropsychiatr. Dis. Treat. 13, 397–406. doi:10.2147/NDT.S123598.

Whitfield-Gabrieli, S., Thermenos, H. W., Milanovic, S., Tsuang, M. T., Faraone, S. V., McCarley, R. W., et al. (2009). Hyperactivity and hyperconnectivity of the default network in schizophrenia and in first-degree relatives of persons with schizophrenia. Proc. Natl. Acad. Sci. U. S. A. 106, 1279–1284. doi:10.1073/pnas.0809141106.

Wood, S. J., Yücel, M., Wellard, R. M., Harrison, B. J., Clarke, K., Fornito, A., et al. (2007). Evidence for neuronal dysfunction in the anterior cingulate of patients with schizophrenia: A proton magnetic resonance spectroscopy study at 3 T. Schizophr. Res. 94, 328–331. doi:10.1016/j.schres.2007.05.008.

Woodward, N. D., Rogers, B., and Heckers, S. (2011). Functional resting-state networks are differentially affected in schizophrenia. Schizophr. Res. 130, 86–93. doi:10.1016/j.schres.2011.03.010.

Yan, H., Tian, L., Yan, J., Sun, W., Liu, Q., Zhang, Y. B., et al. (2012). Functional and Anatomical Connectivity Abnormalities in Cognitive Division of Anterior Cingulate Cortex in Schizophrenia. PLoS One 7. doi:10.1371/journal.pone.0045659.

Yoav Benjamini; Yosef Hochberg (1995). Controlling the False Discovery Rate: A Practical and Powerful Approach to Multiple Testing. R. Stat. Soc. Ser. B (Methodol.) 57, 289–300.

Yu, Q., Du, Y., Chen, J., He, H., Sui, J., Pearlson, G., et al. (2017). Comparing brain graphs in which nodes are regions of interest or independent components: A simulation study. J. Neurosci. Methods 291, 61–68. doi:10.1016/j.jneumeth.2017.08.007.

Zhang, Z., Liu, G., Yao, Z., Zheng, W., Xie, Y., Hu, T., et al. (2018). Changes in dynamics within and between resting-state subnetworks in juvenile myoclonic epilepsy occur at multiple frequency bands. Front. Neurol. 9, 1–11. doi:10.3389/fneur.2018.00448.

Zhi, D., Calhoun, V. D., Lv, L., Ma, X., Ke, Q., Fu, Z., et al. (2018). Aberrant Dynamic Functional Network Connectivity and Graph Properties in Major Depressive Disorder. Front. Psychiatry 9, 1–11. doi:10.3389/fpsyt.2018.00339.

Zou, H., and Hastie, T. (2005). Regularization and variable selection via the elastic net. J. R. Stat. Soc. Ser. B Stat. Methodol. 67, 301–320. doi:10.1111/j.1467-9868.2005.00503.x.

