## Supplementary Information for "Aberrant Dynamic Functional Connectivity of Default Mode Network in Schizophrenia and Links to Symptom Severity"

***Classification and feature selection results for each state***

For the COBRE dataset, the area under the receiver operating characteristic curve (AUC) was 0.72±0.02. 0.66±0.01, 0.76±0.02, 0.71±0.02, and 0.66±0.03 in state 1, state 2, state 3, state 4, and state 5, respectively (Figure 4). The feature learning results showed that $C_3$ (the connectivity between PCu1 and ACC1 or PCu1/ACC1), $C_{10}$ (PCu2/PCC1), $C_{19}$ (ACC2/PCC1) were the most important features in the classification between SZ and HC in state 1 (corrected $p<0.05$). For state 2, the most important features were $C_1$ (PCu1/PCu2), $C_5$ (PCu1/PCC1), $C_7$ (PCu2/PCu3), $C_8$ (PCu2/ACC1), $C_{10}$ (PCu2/PCC1), $C_{12}$ (PCu3/ACC1), $C_{15}$ (PCu3/PCC2), $C_{16}$ (ACC1/ACC2), and $C_{19}$ (ACC2/PCC1) (corrected $p<0.05$). In state 3, $C_7$ (PCu2/PCu3), $C_9$ (PCu2/ACC2), $C_{12}$ (PCu3/ACC1), and $C_{14}$ (PCu3/PCC1) were the most important features in SZ vs. HC classification (corrected $p<0.05$). $C_1$ (PCu1/PCu2), $C_9$ (PCu2/ACC2), $C_{10}$ (PCu2/PCC1), $C_{11}$ (PCu2/PCC2), and $C_{19}$ (ACC2/PCC1) were the most important features in the state 4 classification (corrected $p<0.05$). In state5, $C_8$ (PCu2/ACC1), $C_{11}$ (PCu2/PCC2), $C_{15}$ (PCu3/PCC2), $C_{17}$ (ACC1/PCC1), and $C_{20}$ (ACC2/PCC2) most contributed to the classification between SZ and HC (corrected $p<0.05$).

A similar feature selection method on connectivity classification between SZ subjects and HC was conducted for the FBIRN dataset, as shown in Figure 5. The classification AUC between SZ and HC was 0.72±0.01, 0.68±0.01, 0.67±0.02, 0.69±0.01, and 0.68±0.02 in state 1, state 2, state 3, state 4, and state 5, respectively. $C_1$ (PCu1/PCu2), $C_6$ (PCu1/PCC2), $C_{10}$ (PCu2/PCC1), $C_{14}$ (PCu3/PCC1), $C_{16}$ (ACC1/ACC2), and $C_{20}$ (ACC2/PCC2) were the most important features in the discrimination between HC and SZ in state 1. In state 2, $C_{11}$ (PCu2/PCC2), $C_{15}$ (PCu3/PCC2), $C_{16}$ (ACC1/ACC2), $C_{18}$ (ACC1/PCC2), and $C_{19}$ (ACC2/PCC1) were the most important features in the classification between HC and SZ subjects. In state3, $C_5$ (PCu1/PCC1), $C_9$ (PCu2/ACC2), $C_{12}$ (PCu3/ACC1), $C_{14}$ (PCu3/PCC1), $C_{16}$ (ACC1/ACC2) were the most important features (corrected p<0.05). In state 4, $C_5$ (PCu1/PCC1), $C_8$ (PCu2/ACC1), $C_9$ (PCu2/ACC2), $C_{14}$ (PCu3/PCC1), $C_{16}$ (ACC1/ACC2), $C_{19}$ (ACC2/PCC1) were the most important features in the classification between HC and SZ (corrected p<0.05). In state 5, the most important features were $C_{10}$ (PCu2/PCC1), C13 (PCu3/ACC2), $C_{14}$ (PCu3/PCC1), $C_{16}$ (ACC1/ACC2), $C_{18}$ (ACC1/PCC2), $C_{19}$ (ACC2/PCC1), and $C_{20}$ (ACC2/PCC2).

**Supplemental Table S1**

Demographic and clinical details of FBIRN subjects for each site.

| | | *SZ* | *HC* | *P-value* |
|---|---|---|---|---|
| ***FBIRN Site 1*** | **Number** | 21 | 28 | NA |
| | **Age** | 30.04±8.60 | 34.78±9.40 | 0.1 |
| | **Gender(M/F)** | 17/4 | 21/7 | 0.99 |
| | **PANSS (positive)** | 15.72±5.57 | NA | NA |
| | **PANSS(negative)** | 14.11±3.14 | NA | NA |
| ***FBIRN Site 2*** | **Number** | 12 | 10 | NA |
| | **Age** | 44.91±11.34 | 38.10±9.39 | 0.14 |
| | **Gender(M/F)** | 12/0 | 7/3 | 0.62 |
| | **PANSS (positive)** | 16.90±6.70 | NA | NA |
| | **PANSS negative)** | 17.20±7.39 | NA | NA |
| ***FBIRN Site 3*** | **Number** | 24 | 27 | NA |
| | **Age** | 44.41±11.90 | 42.48±12.56 | 0.57 |
| | **Gender(M/F)** | 19/5 | 21/6 | 0.99 |
| | **PANSS (positive)** | 16.95±4.43 | NA | NA |
| | **PANSS negative)** | 16.87±5.91 | NA | NA |
| ***FBIRN Site 4*** | **Number** | 26 | 26 | NA |
| | **Age** | 36.88±12.82 | 35.23±10.56 | 0.61 |
| | **Gender(M/F)** | 21/5 | 20/6 | 0.99 |
| | **PANSS (positive)** | 14.29±3.77 | NA | NA |
| | **PANSS (negative)** | 13.41±4.64 | NA | NA |
| ***FBIRN Site 5*** | **Number** | 14 | 15 | NA |
| | **Age** | 36.64±10.27 | 37.53±9.76 | 0.81 |
| | **Gender(M/F)** | 9/5 | 10/5 | 0.99 |
| | **PANSS (positive)** | 13.14±4.46 | NA | NA |
| | **PANSS (negative)** | 15±6.28 | NA | NA |
| ***FBIRN Site 6*** | **Number** | 29 | 27 | NA |
| | **Age** | 36.27±11.08 | 34.51±10.85 | 0.87 |
| | **Gender(M/F)** | 17/12 | 16/9 | 0.99 |
| | **PANSS (positive)** | 14.34±4.67 | NA | NA |
| | **PANSS (negative)** | 11.93±4.06 | NA | NA |
| ***FBIRN Site 7*** | **Number** | 25 | 27 | NA |
| | **Age** | 39.56±11.80 | 37.56±10.75 | 0.52 |
| | **Gender(M/F)** | 20/5 | 20/7 | 0.99 |
| | **PANSS (positive)** | 16.12±5.41 | NA | NA |
| | **PANSS (negative)** | 14.04±6.82 | NA | NA |

**Note:** SZ: Schizophrenia; HC: healthy control; PANSS: Positive and Negative Syndrome Scale, M: Male, F: Female, NA: not applicable; all p values have been calculated using two-sample t-test.
